# A mutation-level covariate model for mutational signatures

**DOI:** 10.1101/2022.04.30.490152

**Authors:** Itay Kahane, Mark D.M. Leiserson, Roded Sharan

## Abstract

Mutational processes and their exposures in particular genomes are key to our understanding of how these genomes are shaped. However, current analyses assume that these processes are uniformly active across the genome without accounting for potential covariates such as strand or genomic region that could impact such activities. Here we suggest the first mutation-covariate models that explicitly model the effect of different covariates on the exposures of mutational processes. We apply these models to test the impact of replication strand on these processes and compare them to strand-oblivious models across a range of data sets. Our models capture replication strand specificity, point to signatures affected by it, and score better on held-out data compared to standard models that do not account for mutation-level covariate information.

## 1 Introduction

Cancers are caused by somatic mutations accumulated during the organism’s life [1, 2]. Those mutations, are the result of mutational processes varying from exogenous and endogenous DNA damage to faulty DNA repair and replication [3, 4], and leaving unique mutational signatures [1, 2, 5]. Deciphering these signatures and the genome’s exposure to them are key to understanding how it is shaped by the disease. Such mapping was initially done by non-negative matrix factorization (NMF) and its generalizations [1, 6, 7, 8, 9], or refitting methods that infer the exposures given the signatures [10, 11, 12, 13, 14]. More recent work built on topic models that allow to rigorously attribute likelihood to the data and solve the models’ parameters by maximizing it [15, 16, 17, 18, 19]. One of the advantages of the topic model framework, is that it allows to exploit additional information on the data for improved predictions. For instance, [17] used a generalization of Dirichlet multinomial regression [20] to introduce tumor-level covariates in the context of mutational signature modeling.

The aforementioned methods assume that mutational processes work uniformly across the genome. However, it was previously reported that some mutational processes have strand [21, 22, 23] and region [24] biases. In particular, Signatures 2, 13 and 26 (following COSMIC v2 catalogue [25]) were found to have strong replication strand biases [22].

Here we suggest the first mutation-covariate models that explicitly model the effect of different covariates on the exposures of mutational processes. We apply these models to test the impact of genomic and replication strands on these processes and compare them to strand-oblivious models across a range of data sets.

## 2 Methods

### 2.1 Preliminaries

We follow the common convention and assume that there are *M* = 96 mutation categories (denoting a base substitution and its flanking bases), drawn from *K* known signatures. Our data consists of *T* samples, such that a sample *t* is the set of somatic mutations in the respective tumor whose category and the value of a single binary feature (acting as the mutation-level covariate) are known. The per-sample mutation category count of the two possible feature values are denoted *I*_*t,m*_ and *J*_*t,m*_, respectively. In all following models, we assume that the mutational signatures are known.

The basic model we consider, which can be thought of as the probabilistic analog of NMF, is the multinomial mixture model (MMM) [18]. In MMM, the signatures are modeled as multinomial distributions, such that the probability to draw a mutation *m* from signature *k* is notated by *γ*_*k,m*_. Further, each signature has sample-specific probabilities, aka *exposures*. The model specifies a generative process for mutations, where at first a signature is drawn from an exposure vector. Then, a mutation category is drawn from the signature vector (see Figure 1 for a plate notation). A shortcoming of this model is the fact that exposure vectors of different samples are assumed to be independent.

**Figure 1:**
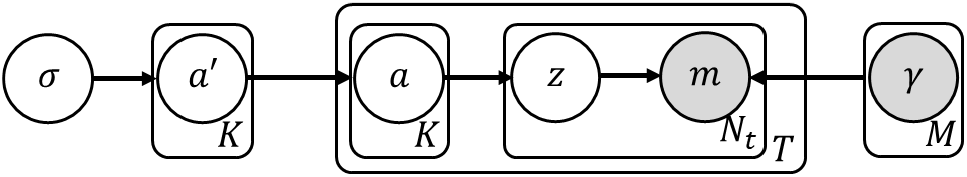
A plate notation of LDA.

To mitigate the latter drawback and generalize to unseen samples, Latent dirichlet allocation (LDA) assumes that the samples share an exposure Dirichlet prior, rather than the exposure vector itself [26]. First, an exposure vector is drawn per-sample from a dirichlet prior. Then, per-mutation, a signature *z* is drawn given the exposure vector and sequentially the mutation category given the signature. Thus, assuming 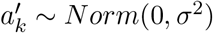 and notating 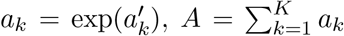 the signature count of sample *t* by *N*_*t,k*_ and 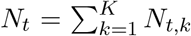 the likelihood of the signature part of the model is given by:

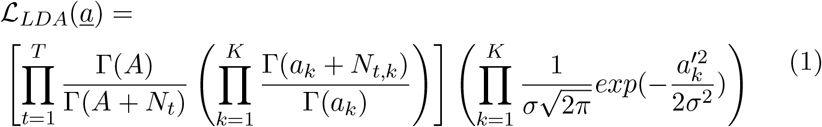

### 2.2 Mutation-level covariate signature model (MCSM)

Similar to Dirichlet multinomial regression, we can generalize the above scheme by dividing it into two parallel LDAs, such that the source LDA of each mutation depends on its feature (see Figure 2 for a plate notation). Using similar notation as before, with *b* in addition to *a* for the second feature and *I* and *J* instead of *N* for the feature-divided data, the likelihood of the signature part of the model is given by:

**Figure 2:**
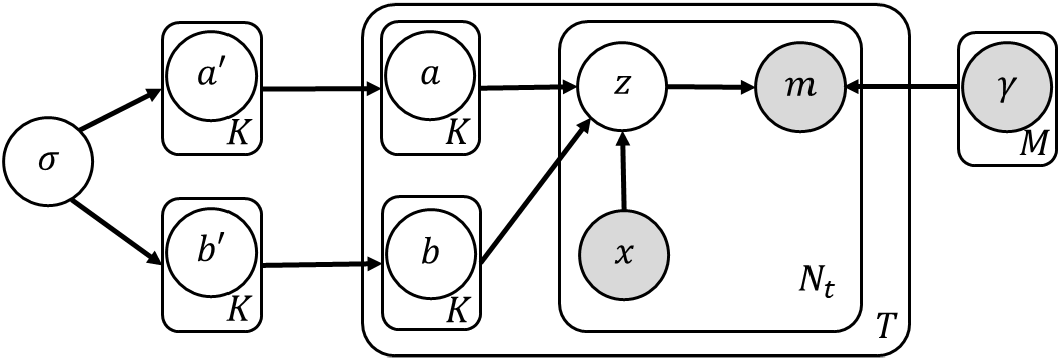
A plate notation of MCSM. Here, *x* is the binary feature that indicates whether*z* is drawn from a Dirichlet distribution with parameter vector *a* or *b*.

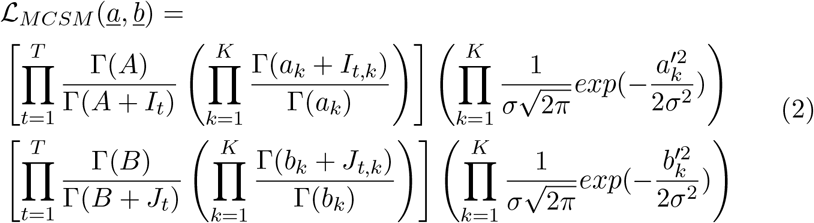

### 2.3 Joint MCSM (JMCSM)

Although the previous MCSM model allows us to integrete mutation level features, it assumes that every tumor has two independent exposure vectors. In reality, since signature exposures are the consequences of genetics and lifestyle, it is reasonable to assume that the two exposures vectors are related rather than independent. To capture this dependency, we assume that a tumor *t* has an *inherent exposure vector* denoted by *e*_*t*_ = (*e*_*t*,1_, …, *e*_*t,K*_) which can be thought of as a tumor level covariate [20, 17] (see Figure 3 for a plate notation). In order to impose that the strand-specific exposure vector is drawn in the proximity of the inherent exposures, we modify the Dirichlet parameters and define them as (*e*_*t*,1_*a*_1_, …, *e*_*t,K*_*a*_*K*_) and (*e*_*t*,1_*b*_1_, …, *e*_*t,K*_*b*_*K*_). The likelihood of the signature part of the model is now given by:

**Figure 3:**
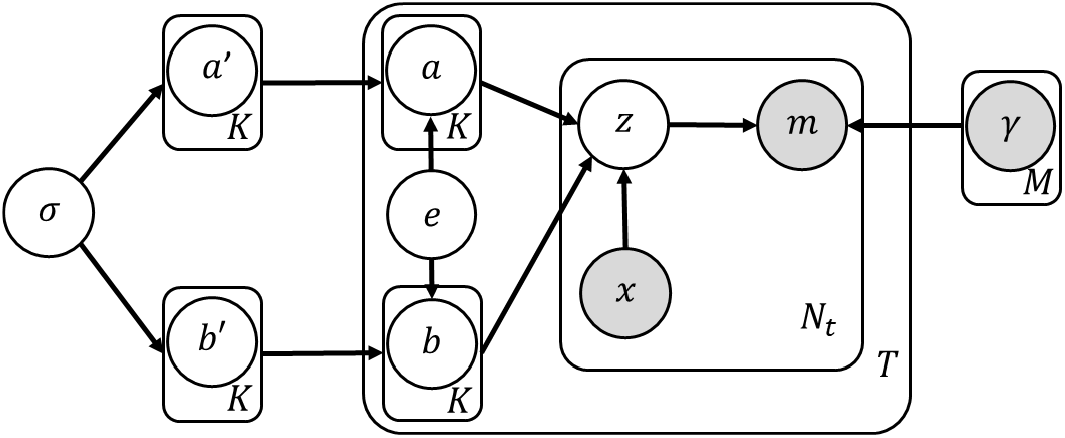
A plate notation of JMCSM.

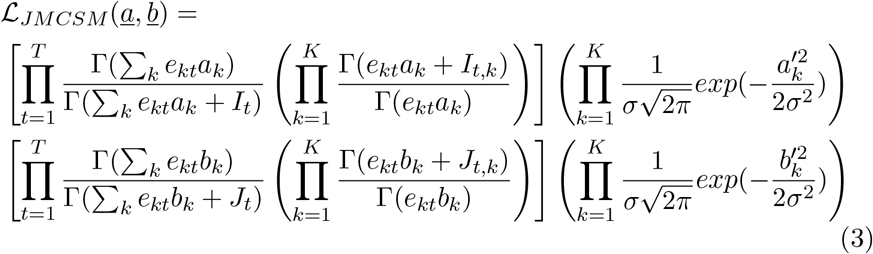

To tease apart the contribution of the joint modeling of strands on model performance, we also define a guided LDA (gLDA) model variant which is fed with external information on the exposures as in JMCSM (see Figure 4 for a plate notation). Its likelihood is given by:

**Figure 4:**
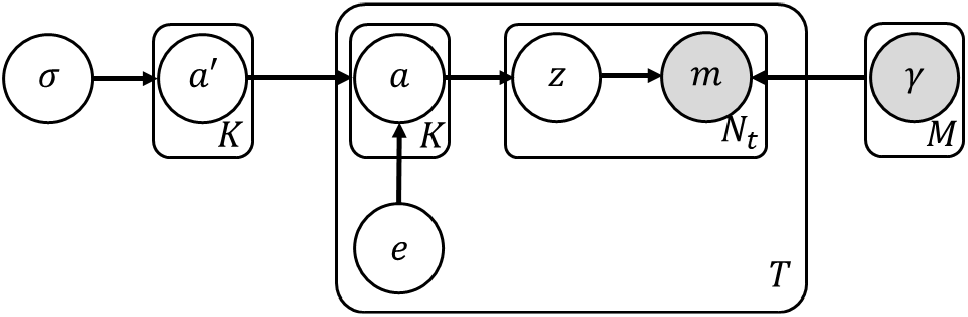
A plate notation of gLDA.

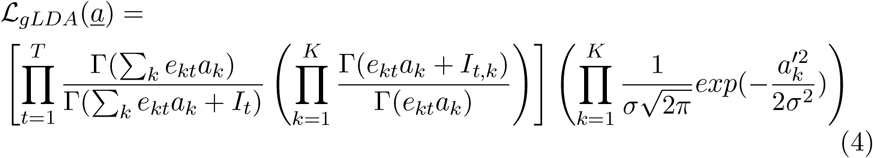

### 2.4 Model Learning

The above topic models (LDA, MCSM, gLDA and JMCSM) can be optimized using stochastic EM (SEM). In SEM, we alternately draw random signature assignment based on the current parameter estimation and then a new set of parameters is estimated given those assignments. As it is not possible to directly calculate the signature assignment probability conditioned on the mutation categories without summing over an exponentially large number of variables, we use Gibbs Sampling to randomly draw the assignments [27]. For instance, we execute Gibbs sampling for JMCSM by iteratively drawing assignments from the conditional probability (for the *n − th* mutation *w*_*n*_ of sample *t*, for the *a* and *I* respetive feature):

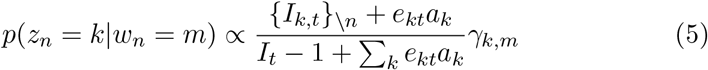

Where { *I*_*k,t*}\*n*_ is the number of mutations drawn with assignment *k*, without counting the current mutation assignment. The process is stopped after a large number of iterations (3,000). Notably, when learning JMCSM we first extract *e*_*t*_ by ignoring the mutation feature and applying the MMM model, optimized via EM. For robustness, we run SEM for 50 iterations and report below the mean held-out empirical likelihood (EL) calculated over the 25 last SEM iterations.

### 2.5 Model evaluation

We evaluate the models by a held-out log-likelihood comparison with a feature oblivious analog: MCSM vs. LDA and JMCSM vs. gLDA. Since the likelihood of LDA and its derivatives is not directly computable, we follow previous works [28], and use the method of EL. In EL we draw a large number of tumor exposures using the estimated parameters, and then calculate the mean log-likelihood of the test set. Specifically, we use *S* = 10, 000 randomizations leading to *S* pairs of expousre vectors for both feature values 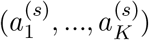 and 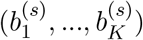, the empirical log-likelihood of MCSM is given by:

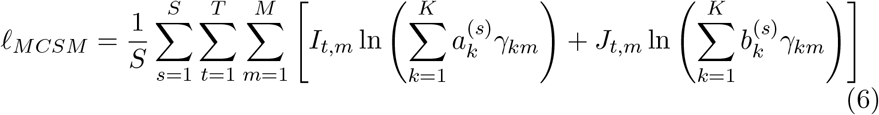

### 2.6 Data Description

To test our novel models we work with Breast Cancer (BRCA), Malignant Lymphoma (MALY) and Chronic Lymphocytic Leukemia (CLLE) from International Cancer Genome Consortiom as in [13] (see Table 1).

**Table 1:** Databases analyzed in this work

| Cancer Type | No. Samples | No. Mutations | Cosmic Signatures |
| --- | --- | --- | --- |
| BRCA | 560 | 3,472,652 | 1, 2, 3, 5, 6, 8, 13, 17, 18, 20, 26, 30 |
| MALY | 100 | 1,220,526 | 1, 2, 5, 9, 13, 17 |
| CLLE | 100 | 270,870 | 1, 2, 5, 9, 13 |

We executed 2-fold cross validation by dividing each sample into two equally-sized subsets. Then, we learned the model parameters using one subset, and calculated the EL using the other one. To extract the inherent exposures, we used MMM on each sample of the train set separately. On JMCSM and gLDA, we then used the inherent exposures as observable variable to learn the model parameters. We also used the pre estimated inherent exposures to calculate the EL. We repeated this scheme twice to evaluate both MCSM and JMCSM compared with their respective comparable model with a single mutation-level covariate: whether the mutation occured on leading or lagging strand. Since we only use mutations on which those feature are relevant and available, the actual usuable size of both database is smaller (see Table 2).

**Table 2:** Number of mutations on each strand

| Cancer Type | Lagging | Leading |
| --- | --- | --- |
| BRCA | 313,219 | 209,488 |
| MALY | 96,177 | 68,365 |
| CLLE | 13,315 | 12,534 |

## 3 Results

We designed novel models for mutational signature analysis that account for mutation-level features. The basic model, MCSM, is a generalization of the standard LDA that learns two different Dirichlet priors for the case of a binary feautre, each corresponding to a different value of the feature. A refined model, JMCSM, accounts also for tumor-level covariates in the form of a vector exposures that ties together the two Dirichlet priors (with an LDA-like benchmark termed gLDA). To test our models we focused on the richest features that we could extract, namely the genomic and replication strand information across different data sets. While Watson/Crick data did not seem to improve learning (see discussion in the Conclusions), notable differences were observed with respect to the lagging/leading strand data (Table 3).

**Table 3:** A comparison between MCSM and LDA, and between JMCSM and gLDA using replication strand as a mutation-level feature. The percentage difference sign indicates whether the strand sensitive model has a better likelihood (negative), and vice versa.

| dataset | MCSM | LDA | diff % | JMCSM | gLDA | diff % |
| --- | --- | --- | --- | --- | --- | --- |
| BRCA | -2,216,362 | -2,215,837 | 0.024 | -1,995,494 | -1,996,589 | -0.055 |
| MALY | -717,961 | -718,491 | -0.074 | -694,699 | -694,937 | -0.034 |
| CLLE | -112,664 | -112,703 | -0.035 | -111,869 | -111,866 | 0.002 |

As evident from the table, the refined JMCSM and gLDA models yield better held-out log-likelihood than their basic versions thanks to their usage of covariate information of *inherent exposures*. While MCSM does not consistently improve upon LDA, JMCSM dominates the other models when tested on the larger datasets, BRCA and MALY.

Next, we wished to pinpoint the signatures with replication strand bias. To this end, we calculated for each signature the log-ratio magnitude of its normalized modification parameters *a*_*k*_/Σ_*i*_*a*_*i*_ and *b*_*k*_/Σ_*i*_*b*_*i*_. As *a* and *b* indicates the relative bias of a signature given a featue, the normalized ratio indicates its intensity. The results are given in Figure 5 for the two larger datasets.

**Figure 5:**
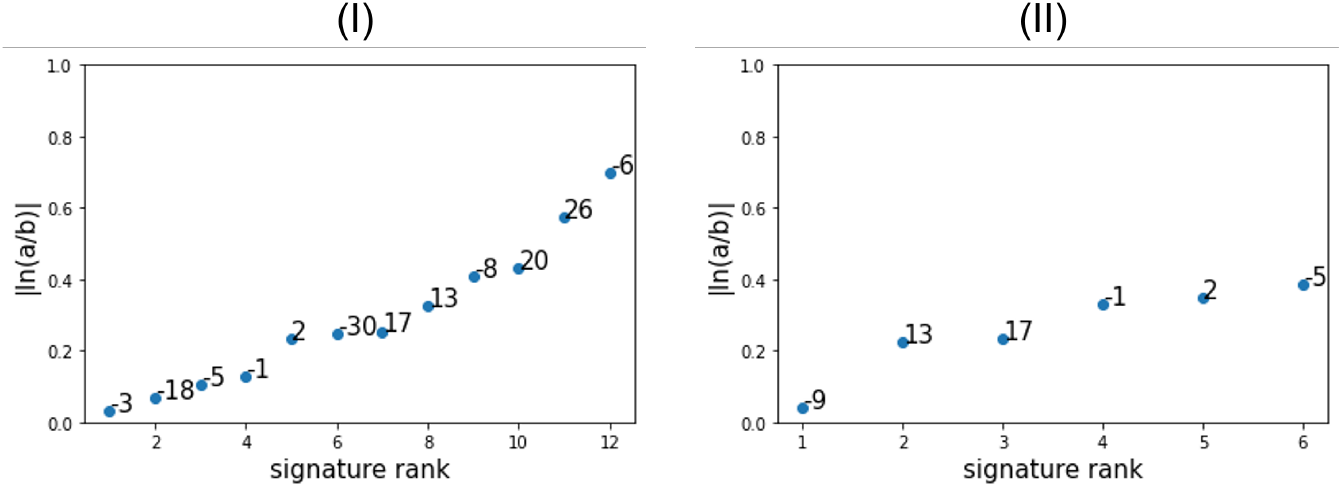
Log-ratio magnitude of *a* and *b* for each signatures in BRCA (I) and MALY (II).

It can be seen that although Signatures 2, 13, 26 are known to have a strong replication strand bias, some other signatures seem to have stronger bias in our framework, most notably Signature 6 which has the highest log-ratio among all tested signatures. A major disadvantage of this method is that it does not take into account the signature frequency in the dataset. A rare signature would have a relatively small effect on the held-out likelihood, and its respective *a* and *b* bias parameters are more prone to overfitting. To tackle this problem, we used a second evaluation of strand bias by calculating the contribution of each signature seperately to the held-out log likelihood. To do so, we compared the likelihood obtained when imposing no bias on the signatures (by averaging the Dirichlet parameters) to the likelihood obtained when allowing only a single signature to be biased (by using its Dirichlet parameter and averaging all others). The results are given for BRCA and MALY in Figure 6.

**Figure 6:**
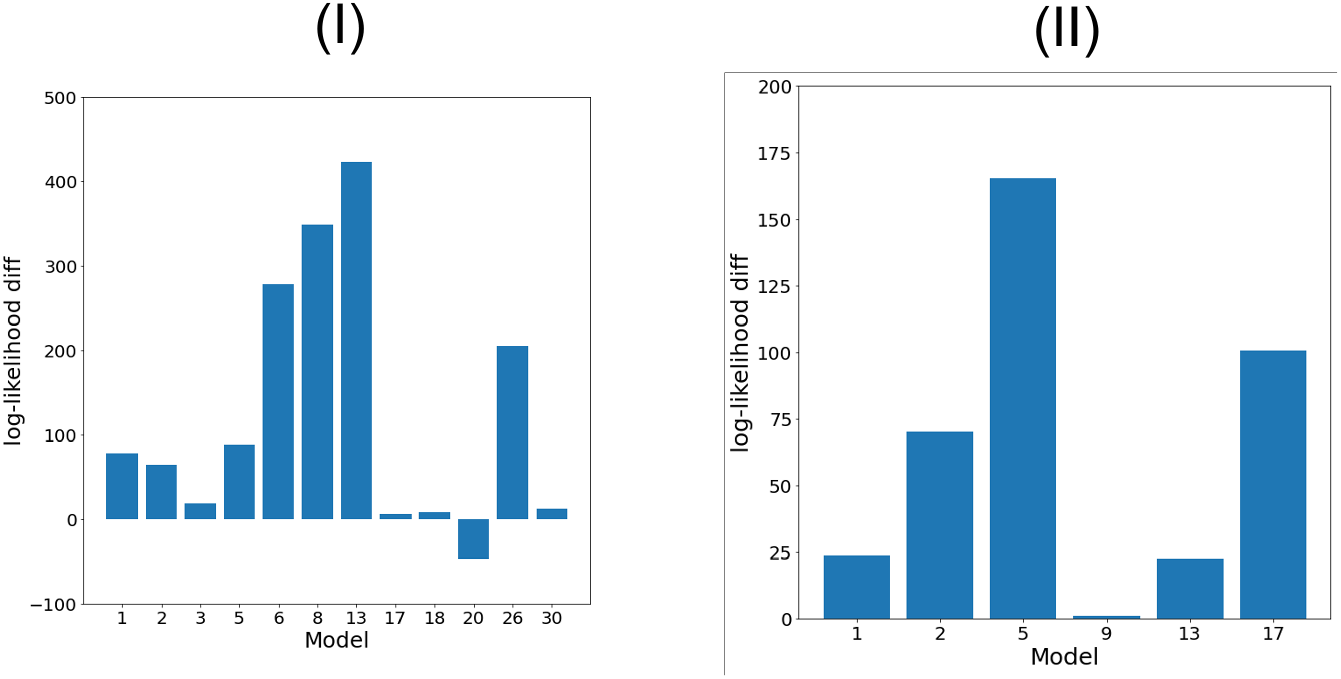
Held-out log-likelihood contribution of each signature for BRCA (I) and MALY (II). The y-axis shows the difference between the log-likelihood of an unbiased model and a single signature bias model. The number under the bars indicates the signature whose bias is maintained.

The results indicate that for BRCA the main sources for the improvement in likelihood are signatures 6, 8, 13 and 26 (the last two are known to have a strong replication strand bias, as stated earlier). Signature 20 (which is also the rarest in BRCA, in terms of exposure averaged over all samples), however, does not seem to display a positive effect on the likelihood contrary to its high bias using the first evaluation. Considering MALY, signatures 2, 5 and 17 are the major likelihood contributers. Signature 13, despite its strong replication strand bias and its impressive contribution in the BRCA case, yields smaller contribution due to the fact that most of the mutations in the database are assigned more likely to other signatures.

## 4 Conclusions

We have developed new topic models to account for mutation-level covariates when learning mutational signatures and their exposures. The models allowed us to pinpoint signatures that display strand bias. Notably, we also applied our models with the genomic strand of the pyrimidine base in the reference sequence as a feature. However, we could detect no consistent advantage of the models over their strand oblivious counterparts, and indeed such biases are not reported in the literature. While we applied our framework in the context of strand information, the models described here can be easily adapted to other binary or categorial features, and could perhaps be used to reveal other biological differences between mutations, for example when considering coding vs. non-coding regions.

## Acknowledgments

This research was supported by a grant from the United States - Israel Binational Science Foundation (BSF), Jerusalem, Israel (to RS and MDML).

## Data and code availability

Code and links to data can be found here: https://github.com/Kitay10/Mutation-level-covariates.

